# Genome-specific histories of divergence and introgression between an allopolyploid unisexual salamander lineage and two sexual species

**DOI:** 10.1101/284950

**Authors:** Robert D. Denton, Ariadna E. Morales, H. Lisle Gibbs

## Abstract

Quantifying genetic introgression between sexual species and polyploid lineages traditionally thought to be asexual is an important step in understanding what factors drive the longevity of putatively asexual groups. However, the presence of multiple distinct subgenomes within a single lineage provides a significant logistical challenge to evaluating the origin of genetic variation in most polyploids. Here, we capitalize on three recent innovations—variation generated from ultraconserved elements (UCEs), bioinformatic techniques for assessing variation in polyploids, and model-based methods for evaluating historical gene flow—to measure the extent and tempo of introgression over the evolutionary history of an allopolyploid lineage of all-female salamanders and two ancestral sexual species. We first analyzed variation from more than a thousand UCEs using a reference mapping method developed for polyploids to infer subgenome specific patterns of variation in the all-female lineage. We then used PHRAPL to choose between sets of historical models that reflected different patterns of introgression and divergence between the genomes of the parental species and the same genomes found within the polyploids. Our analyses support a scenario in which the genomes sampled in unisexuals salamanders were present in the lineage ∼3.4 million years ago, followed by an extended period of divergence from their parental species. Recent secondary introgression has occurred at different times between each sexual species and their representative genomes within the unisexuals during the last 500,000 years. Sustained introgression of sexual genomes into the unisexual lineage has been the defining characteristic of their reproductive mode, but this study provides the first evidence that unisexual genomes have also undergone long periods of divergence without introgression. Unlike other unisexual, sperm-dependent taxa in which introgression is rare, the alternating periods of divergence and introgression between unisexual salamanders and their sexual relatives could reveal the scenarios in which the influx of novel genomic material is favored and potentially explain why these salamanders are among the oldest described unisexual animals.

## Introduction

The maintenance of sexual reproduction is the “queen of problems in evolutionary biology” because the ubiquity of sexual reproduction seems difficult to explain given the evolutionary advantages of asexual reproduction (Bell 1982). A resolution to this paradox is that sexual taxa are more robust to extinction in the evolutionary long-term due to increased genetic variability, whereas asexuals suffer the cost of reduced genetic variation that acts to limit long-term persistence (Lively and Dybdahl 2000; Goddard et al. 2005; Loewe and Lamatsch 2008). Generally this appears to be the case, as lineages of asexual taxa tend to be younger compared to related sexual species (Neiman et al. 2009). Nonetheless, there are some putative “asexual” lineages that are millions of years old, which underlines the need to understand the evolutionary mechanisms that allow these putative asexual lineages to persist over evolutionary time.

One hypothesis for the presence of old “asexual” lineages that has received increasing support is that these lineages persist because cryptic introgression occurs between long-lived asexuals and closely-related sexual species, which mitigates both Muller’s ratchet and any Red Queen effects (Hurst and Peck 1996; Beukeboom and Vrijenhoek 1998). Thus, identifying and quantifying these “signs of sex” in putatively asexual lineages has become an essential part of understanding what mechanisms produce younger lineage ages in most non-sexual taxa (Schurko et al. 2009).

Support for this hypothesis comes from recent studies that show few ancient asexual lineages are in fact completely asexual (Mable 2007; Gladyshev et al. 2008; Boschetti et al. 2012; Sánchez Navarro et al. 2013). Introgression from sexual species is documented in multiple eukaryotes, including unisexual fishes (Vrijenhoek 1994, 1998), Iberian minnows (Alves et al. 2001), European water frogs (Spolsky and Uzzell 1986), nematodes (Lunt 2008), and snails (Neiman and Lively 2005), yet precise estimates of the rate and magnitude of introgression are rare (see Schurko *et al*. 2009 for review). Many introgression examples come from taxa that are gynogenetic, a form of reproduction in which all-female lineages require the mechanical or chemical contribution of sperm in order to initiate egg development, but the embryos do not inherit genetic information from the sperm donor (Dawley and Bogart 1989). The tempo of genome introgression among these gynogenetic lineages with “leaky” reproduction is an important consideration for understanding the mechanisms that determine lineage age. If regular introgression occurs, the prediction is that genome replacement will result in asexual lineages becoming genetically identical to their parental species, which raises questions about the selective advantages of unisexuals over sexuals (Charney 2012). In contrast, if introgression is limited to a specific time period in the past, genetic diversity in asexual lineages might be better explained by recombination or efficient removal of harmful mutations (Loewe and Lamatsch 2008). For example, the unisexual fish *Poecilia formosa* originated through hybridization ∼100 kya, continued a short period of backcrossing with the parental species, but displays little evidence of contemporary gene exchange (da Barbiano et al. 2013). Thus, the high levels of observed genetic diversity in this lineage is best explained by a combination of mitotic recombination and preservation of the initial genetic diversity from backcrossing. Until recently, inferences about the historical timing of ingression in polyploid animals were not possible not only due to a lack of genomic resources, but also the difficulty in distinguishing incomplete lineage sorting from hybridization (Choleva et al. 2014) and the bioinformatics challenges associated with identifying genome-specific variation in polyploids (Roux and Pannell 2015). However, recent methodological and analytic developments now approach these challenges directly.

One lineage where introgression is hypothesized to play a large role in lineage persistence is the all-female salamanders of the genus *Ambystoma*. These animals are the oldest known lineage of unisexual vertebrates (∼5 million years) and reproduce through kleptogenesis, a mode of reproduction in which unisexual female salamanders can occasionally “steal” sperm from the males of congeneric sexual salamanders (Bogart et al. 2007; Bi and Bogart 2010). Unisexual salamanders initiate egg production through stimulation with sperm from one of five sexual *Ambystoma* species, and generally discard the male’s contribution, producing maternal clones. However, the spermataphore’s haploid genome may occasionally be incorporated into offspring, resulting in an increase in ploidy or the potential substitution of a genome originally present in the female (Bi et al. 2008). In addition to their surprisingly old lineage age, unisexual *Ambystoma* are one of the most widespread salamander taxa in North America and appear to be highly successful ecologically, outnumbering sexual *Ambystoma* by as many as 2:1 in specific populations (Bogart and Klemens 2008).

Compared to other “leaky” gynogenetic reproductive systems in which genomic introgression does not occur (Beukeboom and Vrijenhoek 1998; Arioli et al. 2010; Pruvost et al. 2013), the rate of genome contribution from sexual *Ambystoma* species into the unisexual lineage is frequent across generations and short evolutionary time scales (Bogart et al. 1989, 2007; Gibbs and Denton 2016).

Estimates of the rate at which sexual species contribute genomes to unisexual *Ambystoma* have varied, with evidence for extensive contributions (27% of individuals each generation; Bogart *et al*. 1989), similar levels of gene flow to that between geographically proximate sexual populations (∼0.2% introgressed genomes per generation; Gibbs and Denton 2016), or no evidence for introgression at all (Spolsky et al. 1992). Part of the reason for these variable estimates may be that the types of genetic markers used and methods of analysis have varied widely. Early studies relied on a single nuclear (Bi et al. 2008) or mitochondrial (Bogart et al. 2007) marker and used qualitative comparisons of variation present in sexuals and unisexuals, whereas more recent studies have additionaly used data from multiple species-specific microsatellite loci combined with coalescent-based models to infer the history of gene flow for specific subgenomes of unisexual individuals (Gibbs and Denton 2016).

All of the above studies focus on contemporary estimates of introgression and provide no perspective on patterns of introgression over the entire five million year evolutionary history of the unisexual lineage (Bi and Bogart 2010). As a result, it is unknown if introgression has proceeded consistently over the last five million years or if introgression may have happened in bursts associated with shifts in distributions, climate fluctuations, or some other environmental factor. Understanding the timing and extent of these introgression events would help evaluate the possible evolutionary mechanisms responsible for the extreme age of this kleptogenetic lineage. For example, if the rate of contemporary introgression as measured by microsatellites (Gibbs and Denton 2016) is consistent over longer time periods, it suggests that there is selection for the stability of allopatric genomes in unisexual populations in the face of rapid introgression of sympatric genomes. In contrast, if the evolutionary history of unisexual salamanders is marked by periods of divergence from related sexual species, the introgression and “sperm theft” that characterizes this unisexual lineage could be a recent phenomenon perpetuated by distributional shifts, forced sympatry due to habitat loss, or changes in climate regimes. Characterizing the evolutionary history of unisexual salamanders’ nuclear genomes would allow us to understand if their unusual lineage age is the result of periodic bouts of introgression or the surprising maintenance of a single unisexual lineage in spite of frequent introgression.

Recovering single nucleotide polymorphisms from the subgenomes of polyploids is a longstanding sequencing and bioinformatic challenge (Dufresne et al. 2014; Salmon and Ainouche 2015). A primary reason for this challenge is the difficulty in distinguishing between homologous markers between subgenomes (difference between parental genomes) and allelic SNPs (polymorphisms unique to a specific subgenome). The explosion of high-throughput sequencing over the last decade has been trailed by an increase in the number of tools and approaches to meet this challenge (Clevenger et al. 2015). The majority of polyploid SNP calling methods utilize mapping techniques that match reads from polyploid individuals to related reference sequences or genomes (McKenna et al. 2010; Li 2011; Page et al. 2013). However, reference-free approaches are becoming more common, including those that rely on detecting asymmetries among polyploid reads (Zohren et al. 2016) or genotype likelihoods (Blischak et al. 2018).

While these approaches have provided multiple strategies for genotyping the subgenomes of polyploids in general, unisexual *Ambystoma* provide unique challenges. Their large (> 25 gb) and highly repetitive genomes have limited the ability to successfully sequence and assemble salamander genomes (Sun et al. 2012; Keinath et al. 2015; Dodsworth et al. 2016). Genome resources for amphibians are slowly appearing (Sun et al. 2015; Session et al. 2016; Nowoshilow et al. 2018), but the majority of these projects are in frogs that generally have smaller genomes than salamanders (Gregory 2016). Meanwhile, there have been successful efforts to obtain reduced representation libraries for salamanders with large genomes by either modifications of protocols to reduce off-target reads (McCartney-Melstad et al. 2016) or by using standard sequence capture methods (Newman and Austin 2016). For differentiating between the intraspecific nuclear genomes within polyploid *Ambystoma*, loci with highly conserved regions and more variable flanking sequence, such as ultraconserved elements (Faircloth et al. 2012), are especially attractive as a way to potentially provide both species-level and population-level resolution. Even with appropriate genomic resources that can delimit the subgenomes of polyploids and provide adequate resolution for analyses, only recently have techniques been developed that address the challenges of identifying divergence between species in the face of gene flow (Solís-Lemus and Ané 2016; Jackson et al. 2017b). Here, we use a recently developed model selection approach that considers both gene flow and divergence, PHRAPL, to analyze the evolutionary history of UCE loci from polyploid unisexual salamanders that were assigned back to their parental species of origin. Together, this approach provides the first nuclear perspective on the evolutionary history of introgression and divergence in this unusual vertebrate lineage.

## Materials and Methods

### Sample collection and preparation

We collected adults of two sexual species, *A. jeffersonianum* (N = 28) at three sites across Ohio and adult *A. laterale* (N = 17) at one site in southeastern Michigan and one site in northwest Ohio, during spring breeding migrations between 2007 and 2013 (Table 1). Unisexual *Ambystoma* (N = 50) were also collected from all five sites. Two of these sites (Kitty Todd Nature Preserve and Big Creek Park) were the same populations as analyzed by Gibbs and Denton (2016). We removed a small (< 5mm) amount of tail tissue from each adult and then extracted DNA from the collected tissue using Qiagen DNeasy kits (Qiagen, Valencia, CA, USA). Each extraction was checked for DNA quality via agarose gel, and then analyzed in two ways. First, we confirmed the identity of unisexual individuals using mitochondrial DNA (primers F-THR and R-651; McKnight and Shaffer 1997, Bogart et al. 2007) and determined ploidy number and biotype identity using the a SNP assay designed to determine unisexual salamander genome composition (Greenwald and Gibbs 2012).

**Table 1.** Sampling locations for Ambystoma laterale, A. jeffersonianum, and unisexual Ambystoma with genomes from both sexual species as indicated by abbreviations in parenthesis (LJJ = triploid with one A. laterale genome, two A. jeffersonianum genomes).

| Site | Location | County | Group | N |
| --- | --- | --- | --- | --- |
| Big Creek Park | Northeast Ohio | Geauga | unisexual *Ambystoma* (LJJ, LJJJ) | 17 |
|  |  |  | *A. jeffersonianum* | 14 |
| Orondorf Wetlands | Central Ohio | Delaware | unisexual *Ambystoma* (LJJ) | 9 |
|  |  |  | *A. jeffersonianum* | 8 |
| Fort Ancient | Southwest Ohio | Warren | unisexual *Ambystoma* (LJJ, LJJJ) | 8 |
|  |  |  | *A. jeffersonianum* | 6 |
| Kitty Todd Nature Preserve | Northwest Ohio | Lucas | unisexual *Ambystoma* (LLJ) | 8 |
|  |  |  | *A. laterale* | 8 |
| E.S. George Reserve | Southeast Michigan | Livingston | unisexual *Ambystoma* (LLJ) | 8 |
|  |  |  | *A. laterale* | 9 |

Second, we generated UCE loci using the full tetrapod UCE probe set (5,472 probes: http://ultraconserved.org/; Faircloth et al. 2012), which successfully isolate UCE loci from other salamander species (Newman and Austin 2016). Samples at concentrations of 35-50 ng/µL at 35 µL volumes were sent to RAPiD Genomics (Gainesville, FL), where sample libraries were prepared and enriched using standard Illumina TruSeq adapters with an 8 bp index and the the full tetrapod UCE probe set. Prepared libraries where then sequenced using a full lane of an Illumina HiSeq machine with 100bp paired end reads.

### Ultraconserved element processing

Demultiplexed reads were first inspected for quality with FASTQC (v. 0.11.5; Andrews 2010). We then filtered reads by removing adapter sequences and low quality or ambiguous reads using Trimmomatic (Bolger et al. 2014) as implemented in Illumiprocessor (v. 2.0.7; Faircloth 2013). For the two sexual species only, we then assembled reads *de novo* using Trinity with default parameters (v. 2.0.6; Grabherr et al. 2013) as implemented in Phyluce (v. 1.5; Faircloth 2016). We then used Phyluce to extract contigs that were enriched with UCE probe sequences. At this point, we removed three *A. jeffersonianum* individuals that had few loci successfully extracted when compared to the remaining samples.

### Building reference sequences and calling polyploid genotypes

To extract UCE loci from either the *A. laterale* or *A. jeffersonianum* genomes within each unisexual individual, we first built pseudo-references of extracted UCE sequences and flanking regions for each sexual species. For both species, we chose the individual with the highest number of base pairs per read from each site (two *A. laterale* individuals, three *A. jeffersonianum* individuals). We then assembled the post-Illumiprocessor trimmed reads for these individuals using Trinity, creating a reference set of contigs for each species that represents all the sites where that species was sampled. We then used modified scripts from Harvey et al. (2016) to extract contigs that map to UCE probes and account for those mapped to multiple loci.

We used the above consensus reference sequences for each sexual species to identify single nucleotide polymorphisms (SNPs) within the unisexual individuals from either *A. laterale* or *A. jeffersonianum* subgenomes using SWEEP (Clevenger and Ozias-Akins 2015). This methodology was developed and validated for distinguishing SNPs between similar subgenomes of polyploid plants, and is based on using subgenome polymorphisms as anchors to assign true SNPs between genotypes. First, BWA-mem (Li 2013) was used to map cleaned reads to each sexual reference sequence for each subgenome (L or J). This process created two read pileups for each unisexual individual, one representing those reads mapped to *A. laterale* and one representing *A. jeffersonianum*. The sorted, indexed BAM files for all unisexual samples that map to one of the reference species were then input into SWEEP along with the indexed reference sequence using a minimum ratio of alternate allele to reference allele of 4, low stringency, and a sliding window size of 100 (the same as read lengths). The variant call file produced by the SWEEP pipeline was then separated into a single file for each individual and non-variant sites in each sequence were replaced with the reference base call using the *SelectVariants* and *FastaAlternateReferenceMaker* tools in the Genome Analysis Toolkit (GATK; McKenna et al. 2010, DePristo et al. 2011). Although the SWEEP pipeline may have identified multiple SNPs associated with different subgenomes, this information was condensed into a single sequence by randomly choosing an alternate SNP compared to the reference. This decision was made due to the difficulty in phasing the SNPs correctly across multiple subgenomes from a single parent. Thus, the sequence information from the unisexual individuals represents a sample of their *A. laterale* (unisexual-L) or *A. jeffersonianum* (unisexual-J) genomes regardless of the number of those subgenomes. Finally, each locus was aligned using MAFFT (v. 7.222; Katoh and Standley 2013) as implemented in Geneious (v. 10; Drummond et al. 2012). Alignments with less than two individuals from either the sexual species or the unisexuals were removed.

### Hierarchical model selection

PHRAPL uses gene trees from multiple loci as input to estimate and compare the statistical fit of demographic models. We first built input gene trees using two separate data sets, *A. jeffersonianum* genomes (*A. jeffersonianum* individuals and unisexual-J) and *A. laterale* genomes (*A. laterale* individuals and unisexual-L). For each locus, we followed Jackson et al. (2016) and generated gene trees with RAxML (v. 8.2.7; Stamatakis 2006) using Rapid hill-climbing and a GTR-GAMMA model of evolution.

We assessed the fit of five initial models to test if 1) genomes from unisexuals had indistinguishable evolutionary histories from the sexual species and 2) if there is or has been gene flow between lineages following divergence (Figure 1). To provide greater resolution on the timing of introgression for the best-supported model for both species-unisexual comparisons, we generated a second model set that included only models representing constant introgression, secondary contact, or divergence with gene flow. We set the divergence time to the value estimated in the most supported model from model set one above, and used the estimated nuclear divergence date for *A. laterale* and *A. jeffersonianum* from Pyron (2014; 12.4 mya) to scale the divergence time estimates of our analysis. We then tested models of secondary contact and divergence with gene flow across six, ∼500,000 year time slices (∼500,000-3.4 mya years; Figure 2A). Finally, to investigate the resolution of our data within the most recent time period, we tested a final set of models at finer time scales (intervals of 50,000 years from ∼50,000-500,000 years ago; Figure 2).

**Figure 1.**
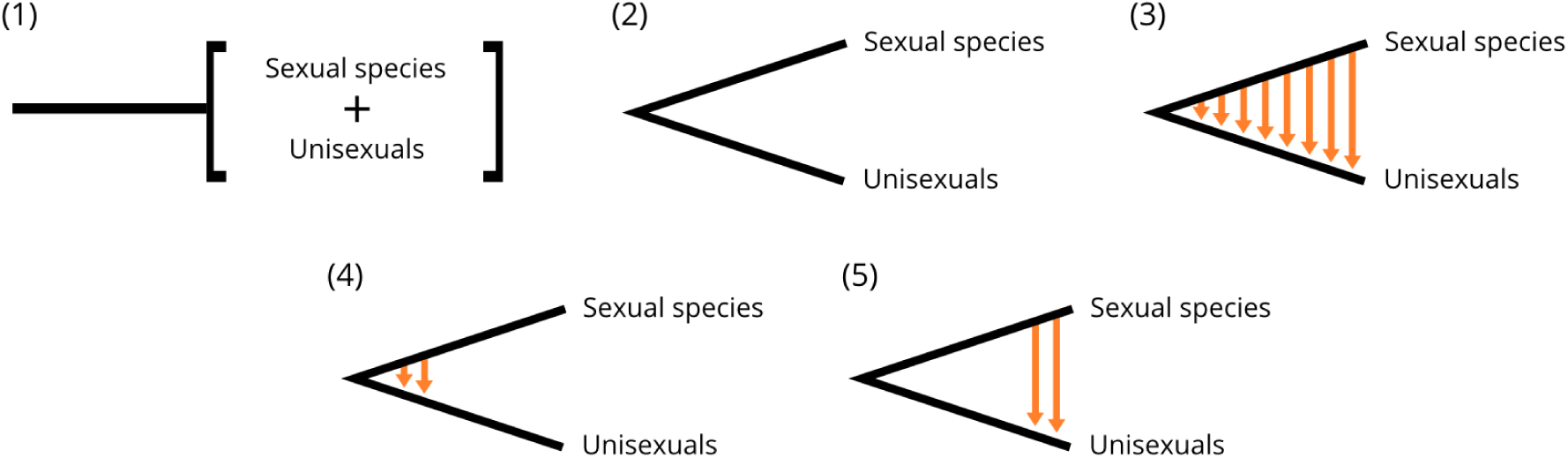
Visual description of initial model set used for PHRAPL model selection. Sexual species can be either *A. laterale* or *A. jeffersonianum*, with unisexuals corresponding to the subgenomes present in the unisexuals that were captured from either sexual species. This initial model set includes a representation of frequent introgression that results in a single lineage (1), divergence of sexual genomes after initial introduction into the unisexual lineage (2), and three broad models that incorporate the asymmetric genomic introgression from either sexual species into the unisexual lineage (3-5).

**Figure 2.**
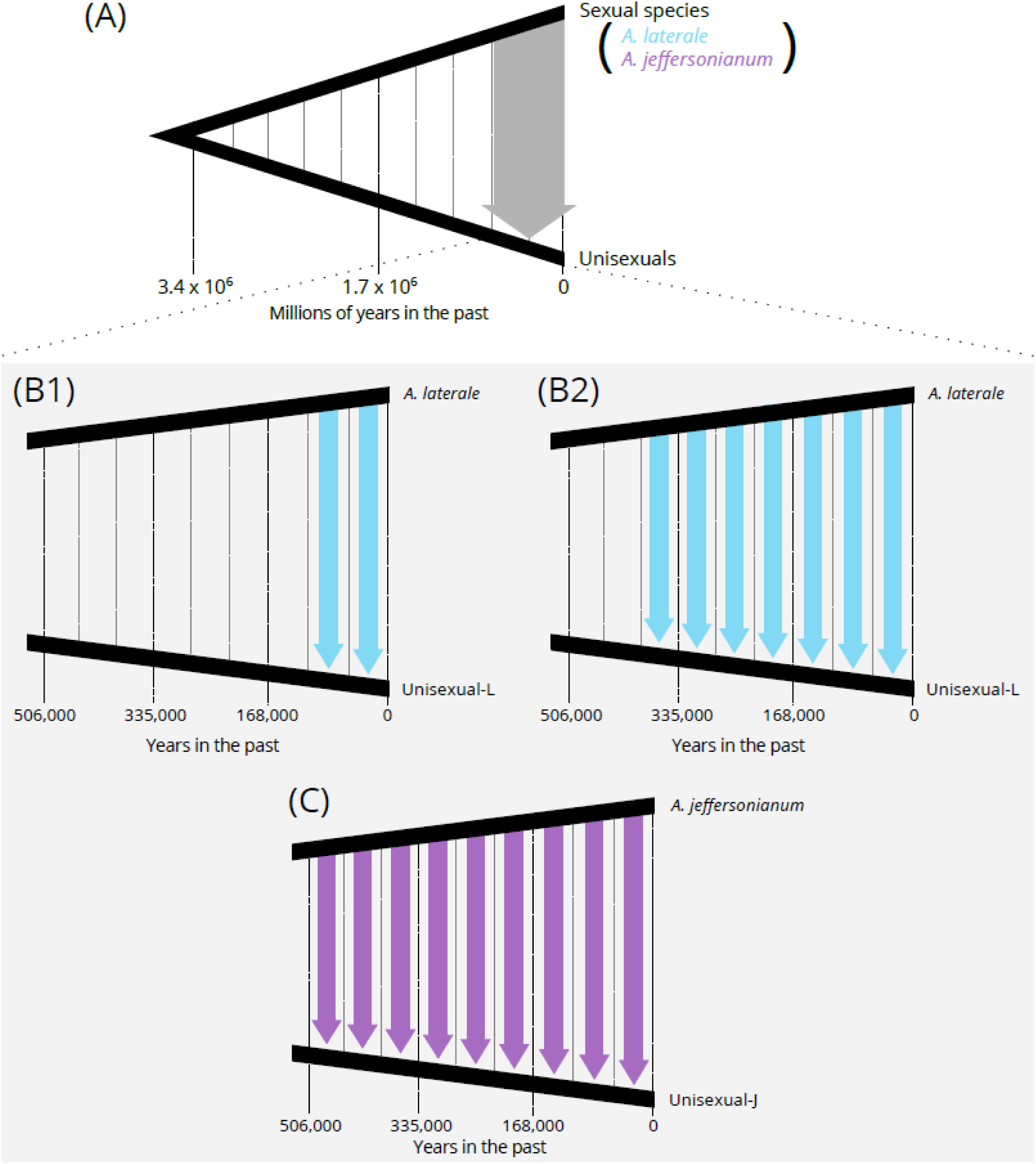
Results of PHRAPL model selection for second set of models for both *A. jeffersonianum* and *A. laterale* and their representative genomes within the unisexual salamander lineage (A). For both, a model of secondary contact is most supported, with divergence taking place approximately 3.4 million years in the past and gene flow beginning in the last ∼563,000 years. Panels B and C visualize the best supported models of the third model set for *A. laterale*, *A. jeffersonianum*, and representative genomes within the unisexual lineage over 50,000-year time slices within the last ∼500,000 years. Models B1 (wAIC = 0.72) and B2 (wAIC = 0.25) for *A. laterale* and representative genomes within the unisexual lineage describe introgression beginning ∼113,000 and ∼397,000 years ago, respectively. Panel C visualizes the most supported models of the third model set for *A. jeffersonianum* and representative genomes within the unisexual lineage, where introgression begins between 507,000 (wAIC = 0.60) and 451,000 (wAIC = 0.33) years in the past. All divergence estimates are calculated as number of generations by PHRAPL and anchored to the estimated divergence time between *A. laterale* and *A. jeffersonianum* (12.4 mya; Pyron 2014).

We performed model selection as implemented in PHRAPL (Jackson et al. 2017a,b) using the input gene trees from each dataset separately (*A. laterale* and unisexual-L, *A. jeffersonianum* and unisexual-J). Gene trees were subsampled at random with replacement 100 times, sampling two individuals per lineage in each replicate. We conducted a simulation of 100,000 gene trees using a grid of parameter values for divergence time (*t*) and migration (*m*) designed to encompass the range of potential values in each dataset (*t* = 0.30, 0.58, 1.11, 2.12, 4.07, 7.81 and *m* = 0.10, 0.22, 0.46, 1.00, 2.15, 4.64).

The lnL and Akaike’s information criterion (AIC) of each model were calculated based on the proportion of matches between simulated and empirical trees. Akaike weights (wAIC) were used to compare models and calculate metrics analogous to model probabilities from 0 (low support) to 1 (high support).

## Results

### Capture probes and polyploid genotyping

Sequencing of the UCE-enriched libraries produced a total of over 422 million reads (mean per individual ± SE = 4.4 x 10^6^ ± 218,508), and these reads were assembled into more than 1,500 contigs per individual. We extracted 2,834 UCE loci from the *A. laterale* and *A. jeffersonianum* individuals combined with a mean length of 331 base pairs. Following the mapping, aligning, and filtering of the unisexual reads, we completed further analyses using 1,183 loci in the *A. laterale* dataset (*A. laterale* and uni-L) and 1,203 loci in the *A. jeffersonianum* dataset (*A. jeffersonianum* and unisexual-J). These final loci were 606 base pairs long on average (range = 224-988). The number of variable sites per locus was higher in *A. jeffersonianum* loci (mean = 3.8) compared to *A. laterale* loci (mean = 1.8), and the coverage for each locus set was similar between *A. jeffersonianum* loci (mean =53.8, range = 22.8-61) and *A. laterale* loci (mean = 51.6, range = 26.5-61.6).

### Hierarchical model selection

The PHRAPL analysis found strongest support for a model of secondary contact (recent gene flow following divergence approximately 3.4 mya) for both datasets (Figure 1, model 5). In the case of *A. jeffersonianum* individuals and unisexual-J loci, a model of secondary contact had a wAIC of 0.66, with no other model with a wAIC of more than 0.19 (Supplementary Table S1). The *A. laterale* dataset was less definitive, with a wAIC of 0.55 for a model of continuous secondary contact and a wAIC of 0.37 for divergence with gene flow. For either *A. laterale* or *A. jeffersonianum* data sets, models that represent either divergence without gene flow or no evidence of divergence were not supported (wAICs << 0.001). Together, these initial models provided support for introgression from each sexual species into the unisexual individuals (Bogart et al. 2007; Gibbs and Denton 2016). However, this introgression does not happen frequently enough to support a scenario where either species and the respective genomes within the unisexual lineage form a single genetically-homogeneous lineage, nor a scenario where introgression happens rarely enough to support a single introgression and subsequent diversification.

For models with greater resolution on the timing of introgression (i.e. with 500,000 year time slices), analyses of both data sets showed the highest support for models of secondary contact (Supplementary Table S1), and each data set best supported a model where contact started within the last 500,000 years (Figure 2A; wAIC = 0.93 for *A. jeffersonianum* loci and 0.97 for *A. laterale* loci). For the final set of *A. laterale* models with intervals of 50,000 years, two had comparatively higher support that the others: one in which introgression began ∼113,000 years ago (wAIC = 0.72) and one in which introgression began ∼397,000 years ago (wAIC = 0.25). For the final set of *A. jeffersonianum* models over the same time period, two models with the greatest support suggest a scenario of sequential episodes of introgression with one episode beginning ∼507,000 years ago (wAIC = 0.60) and another episode occurring ∼451,000 years ago (wAIC = 0.33). To confirm that there was no support for introgression between *A. jeffersonianum* and unisexuals before ∼507,000 years in the past, we reran this final model set with an additional ∼50,000 time slice (∼507,000-563,000 years in the past). Introgression during the oldest time period was not supported compared to the periods between ∼451,000-507,000 years ago.

## Discussion

The subgenomes of unisexual salamanders that are captured from sexual *A. laterale* and *A. jeffersonianum* show a history of secondary contact, where introgression from each sexual species into the unisexual lineage has not been constant following the origin of the unisexual lineage. However, the timing of the onset of introgression following divergence differs between genomes from each of the two species, implying that they have distinct evolutionary histories. Whereas *A. jeffersonianum* genomes have been consistently introgressed into the unisexuals for the previous ∼500,000 thousand years, the loci from *A. laterale* provide evidence for two supported models that describe more distinct periods of introgression (∼113,000 or ∼397,000 years before present). This analysis is the first to provide evidence for subgenome-specific evolutionary histories for the unisexual salamander system and reveals that the genome introgression associated with kleptogenesis is not a constant process over these time scales but has likely varied over evolutionary time.

### Extracting variation from polyploid subgenomes

A technical advance of this work is developing a process to successfully extract subgenome-specific information from allopolyploid unisexual salamanders. Previously, the ability to distinguish genomic variation between the different parental genomes captured in the unisexual lineage was limited to microsatellites loci that amplify in one species or the other or at different size ranges. While microsatellites have been an important step for making inferences about the movement of genomic variation from sexual species into the unisexual lineage at contemporary timescales (Gibbs and Denton 2016) and population genetics of unisexuals (Denton et al. 2017), microsatellites’ rapid rate of evolution precludes using them to make evolutionary inferences that match the same time scale of the unisexual salamander lineage’s mitochondrial formation (Bi and Bogart 2010). In contrast, the combination of rigidly conserved sequences with more variable flanking regions inherent in UCEs can provide both shallow (Harvey et al. 2016) and deep (Faircloth et al. 2012) phylogenetic utility. Because of the evolutionary distance between the two sexual species most represented in the unisexual salamanders, ∼12.4 million years between *A. laterale* and *A. jeffersonianum* (Robertson *et al*. 2006; Pyron 2014), UCEs were an appropriate set of markers to probe nuclear variation within the past six million years. This process identified loci that were shared between sexual species (N = 622), shared only between a sexual species and their representative genomes in the unisexual lineage (N=758 and 707 for *A. laterale* and *A. jeffersonianum*, respectively), and had levels of polymorphism that were suitable for modeling the evolutionary history of these salamanders. Finally, these loci were generated using the standard probe set for vertebrates, providing another example for feasibility of generating of capture data for amphibians with large, highly-repetitive genomes (Newman and Austin 2016).

A limitation to using the combination of UCEs and the SWEEP genotyping in allopolyploid unisexuals is the inability to distinguish allelic SNPs between similar subgenomes. For example, a given UCE locus for an individual with three *A. jeffersonianum* genomes may display two variants compared to the reference, but it is unknown how these variants are distributed across the three *A. jeffersonianum*-like subgenomes. This prevents the identification of subgenome variation at a haploid level unless the genome in question is only present as a single copy in a unisexual individual (e.g. the single *A. laterale* genome in a LJJ unisexual). The consequence of this limitation is the inability to provide genome-specific estimates of introgression into the unisexual lineage because we cannot quantify the variation at the level of the individual subgenome. Our approach of randomly choosing variants as representative of polymorphism means a loss of information, but we assume that our approach means that the polymorphism observed is an unbiased subset of all variants and should provide an accurate albeit reduced sample of polymorphism that should result in accurate parameter estimates. Further advances in the phasing of polyploid sequences (Krasileva et al. 2013) could solve this problem in the future, but our results currently provide the highest temporal resolution available in this system.

### Kleptogenesis through secondary contact

The rarity of clonal animals suggests that there are limited conditions under which genomes from divergent species can interact in a way that initiates a clonal lineage (Warren et al. 2018). Other all-female vertebrate groups, such as Amazon mollies and whiptail lizards in the genus *Aspidoscelis*, have originated multiple times based on rare hybridization events. However, the signal of backcrossing with parental species is common in both groups (Fujita and Moritz 2010; da Barbiano et al. 2013; Warren et al. 2018). These lineages are interesting for how they have maintained an adequate amount of genetic variation to avoid going extinct from some combination of mutation accumulation and Red Queen dynamics (Howard and Lively 1998; Chou and Leu 2015). The unisexual salamander lineage provides an additional level of complexity to this common scenario among asexual animals. While the nuclear genomes present in unisexual salamanders have only recently began to introgress from sexual species, the nuclear genomes in unisexuals have also been previously isolated for longer than any other clonal vertebrate has been extant. This both calls into question how genomic variation had been maintained for millions of years in these salamanders, but also under what conditions introgression has been reestablished.

Other evidence suggests that the introgression from *A. laterale* and *A. jeffersonianum* into the unisexual lineage is not only recent, but frequent. Previous estimates of gene flow from sexual *Ambystoma* species into the unisexual lineage suggest that rates of asymmetrical gene flow can be nearly as high as those between nearby populations of a bisexual species (Gibbs and Denton 2016). While frequent gene flow may aid in preventing the buildup of deleterious mutations in unisexual salamanders, the rate of introgression estimated by Gibbs and Denton (2016) is high enough to potentially homogenize unisexuals with their sympatric sexual species and create a nuclear clone of the local sexual species (e.g. a diploid unisexual with two genomes from *A. jeffersonianum*; Charney 2012). This argues that if genome introgression is frequent, there must be a mechanism that prevents the addition or substitution of new genomes in order to maintain locally-adapted unisexual biotypes. One hypothesis for the mechanism that underlies the selection for retention of allopatric genomes (e.g. an *A. laterale* genome in a unisexual population allopatric to *A. laterale*) is that these genomes contribute to the maintained coexistence between unisexuals and their sexual hosts by allowing for phenotypic diversification that could reduce competition (McIntyre 2012; Ficetola and Stöck 2016). Support for this hypothesis comes from the observation of balanced gene expression among the subgenomes of a unisexual salamander, indicating that all subgenomes contribute relatively equally to one another regardless of which genome matches the resident sperm donor (McElroy et al. 2017).

The secondary contact models that best account for the patterns of shared variation between unisexual L and J subgenomes and their respective parental species indicate that there were extended periods of time (more than 2 million years) with no introgression from either sexual species into the unisexual lineage and subsequent divergence between those genomes in the parental populations and those trapped in the unisexual lineage. This is surprising because—while there are accounts of salamander communities in which male individuals are undetected and suggestions that unisexual *Ambystoma* can potentially be truly parthenogenetic under some circumstances (Uzzell 1969; Noël et al. 2011) —there is strong evidence that unisexuals cannot produce viable eggs without sperm from a compatible sexual species (*reviewed in* Bogart et al. 2017). Further, in light of the strong signals of introgression from microsatellite data (Gibbs and Denton 2016), it seems unlikely that no introgression occurred over the course of many generations. In contrast—and consistent with our results—is the fact that there are multiple other unisexual lineages that arose via hybridization and show no signs of paternal introgression for hundreds of thousands of years across both gynogenetic (Lampert and Schartl 2008; Stöck et al. 2010) and parthenogenetic (Schön and Martens 2003; Kearney et al. 2006) lineages. Alternatively, the five million year divergence estimate from mitochondrial data may not reflect the true initial introgression of either *A. jeffersonianum* or *A. laterale*. This date represents the putative hybridization between an *A. barbouri-*like maternal ancestor and what is assumed to be a male *A. laterale* (Robertson et al. 2006). However, the specific timing of nuclear introgression for either *A. laterale* or *A. jeffersonianum* could have occurred later in the evolutionary history of the unisexual lineage, potentially exaggerating the timing of divergence in our results. Broadly speaking, the weight of support for secondary contact models overall make it likely that the UCE data supports some degree of divergence between the sexual genomes trapped within the unisexual lineage and those that remain in the sexual species and this scenario needs to be considered when developing adaptive and historical explanations for the evolutionary history of genomes found in *Ambystoma* unisexual salamanders.

### Subgenome specific timeframes of genome introgression

Independent of the secondary contact models, the contrast between the timing of genome introgression from *A. laterale* and *A. jeffersonianum* provides evidence for genome-specific differences in the temporal dynamics of kleptogenesis. Our results suggest continuous introgression from *A. jeffersonianum* from ∼507,000 years in the past to present, while introgression from *A. laterale* is intermittent, occurring in two separate periods (∼113,000 years or ∼397,000 years) before the present. At the least, this supports relative differences in the timing of introgression between *A. laterale* and *A. jeffersonianum*. For the populations sampled here (across Ohio and southeast Michigan), introgression from *A. jeffersonianum* has proceeded continuously through time, while introgression from *A. laterale* may have taken place in what we interpret as two recent, separate bursts.

Two factors could explain differential introgression between sexual species. First, there could be differences in the impact of proximate factors that determine if a nuclear genome from a sexual species is incorporated into unisexual lineage. For example, this process can be affected by temperature in unisexual salamanders (Bogart et al. 1989) and polyploid fish (Itono et al. 2007), where a temperature threshold raises the likelihood of sperm incorporation and a subsequent increase in ploidy. Increasing the water temperature at breeding from 6° C to 16° C can quadruple the number of offspring with additional paternal genomes from either *A. laterale* or *A. tigrinum* (Bogart et al. 1989). These temperatures were determined based on average spring pond temperatures in southern Ontario, where *A. laterale* is the most common sexual species. As *A. laterale* is the most northerly distributed caudate in North America, adaptation to breeding in lower temperatures may have resulted in a lower rate of sperm incorporation into the unisexual lineage even where they remain in sympatry for long periods. Secondly, if the cytological ability to incorporate sperm remains relatively constant between sexual species, historical differences in introgression could also be explained by shifting distributions and long periods of allopatry or sympatry with certain sexual sperm donors. How gynogenetic lineages and the sexual species that they interact with remain in sympatry without gynogens outcompeting sexuals remains an active question (Schlupp 2005; Vergilino et al. 2016), and spatial dynamics are important for understanding the maintenance between sexual and asexual organisms in general (Tilquin and Kokko 2016). Given that sperm-dependent parthenogens can slow the range expansion of their sexual hosts (Janko and Eisner 2009), unisexuals interacting disproportionately with *A. jeffersonianum* may contribute to why *A. jeffersonianum* has not displayed a similar post-glacial range expansion to *A. laterale* (Demastes et al. 2007). However, there is currently a lack of empirical evidence for antagonistic interactions or niche competition between unisexual salamanders and their sexual relatives (Brodman and Krouse 2007; Greenwald et al. 2016).

### Predicting the molecular consequences of differential introgression

Preventing rapid nuclear genome turnover in the unisexual salamanders may lessen the potential negative effects of intergenomic conflict. The evolutionary “mismatch” between the mitochondrial genomes within the unisexual lineage and the nuclear genomes that are putatively adapted to the mitochondrial genomes of the sexual species may provide challenges for unisexuals, especially when considering those protein complexes that are encoded by both mitochondrial and nuclear genomes (*reviewed in* Lane 2011). These protein complexes are central to energy production in eukaryotic cells, and mitonuclear mismatch can reduce the efficiency of ATP production (Harrison and Burton 2006), cause oxidative stress (Monaghan et al. 2009), and result in general physiological limitations (Wolff et al. 2014). If the unisexual lineage has had longer history of introgression with *A. jeffersonianum*, cytonuclear interactions may cause fewer negative effects because of the greater available time for the unisexual mitochondrial haplotypes and new *A. jeffersonianum* nuclear genomes to co-evolve. At the same time, genomes from a particular species that are stranded in the unisexual lineage without sexual rescue are more likely to accumulate deleterious mutations (Loewe and Lamatsch 2008; Hollister et al. 2014). Resolving the balance between cytonuclear evolution and the accumulation of deleterious mutations could be a critical step in explaining the exceptional longevity of the unisexual salamander lineage. We show that unisexual subgenomes can display different evolutionary histories, promoting further comparisons between populations with different histories of sympatry and allopatry in order to test how the timing of introgression influences how unisexual genomes change depending on their genomic composition, environment, and ecological interactions.

## Acknowledgements

We thank L Blyth, J David, K Greenwald, G Lipps, T Matson, and R Muehlheim for providing the tissue samples for this work. We thank J Diaz and C Ries for assistance in the laboratory. We acknowledge M Broe, J Clevenger, J Wendel, H Guanjing, C Grover, C Newman, B Faircloth, and B Arnold for their technical advice. This work was funded by a National Science Foundation Doctoral Dissertation Improvement Grant (#1600655) awarded to RDD and HLG.

---

**Supplementary Table S1.**
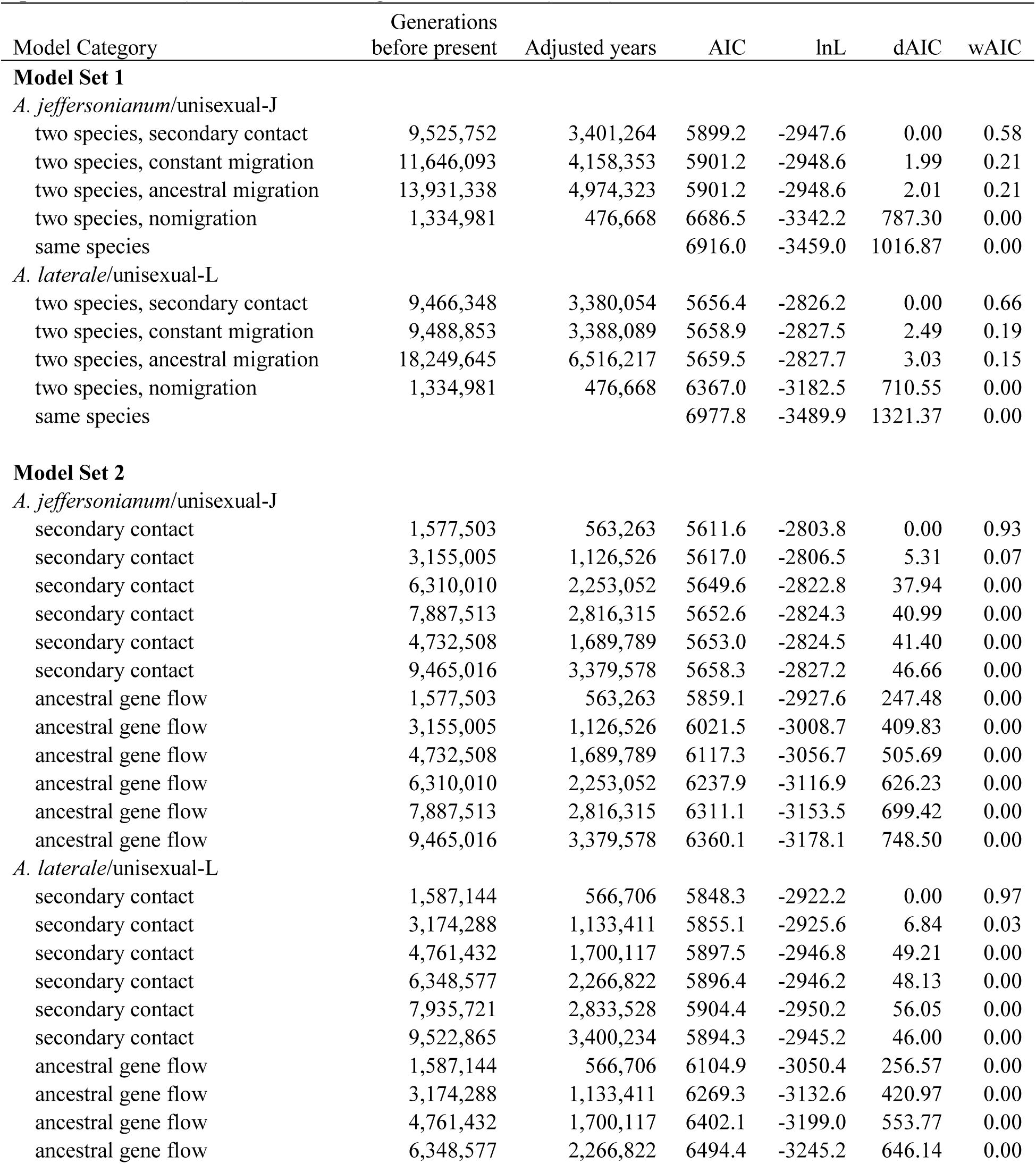
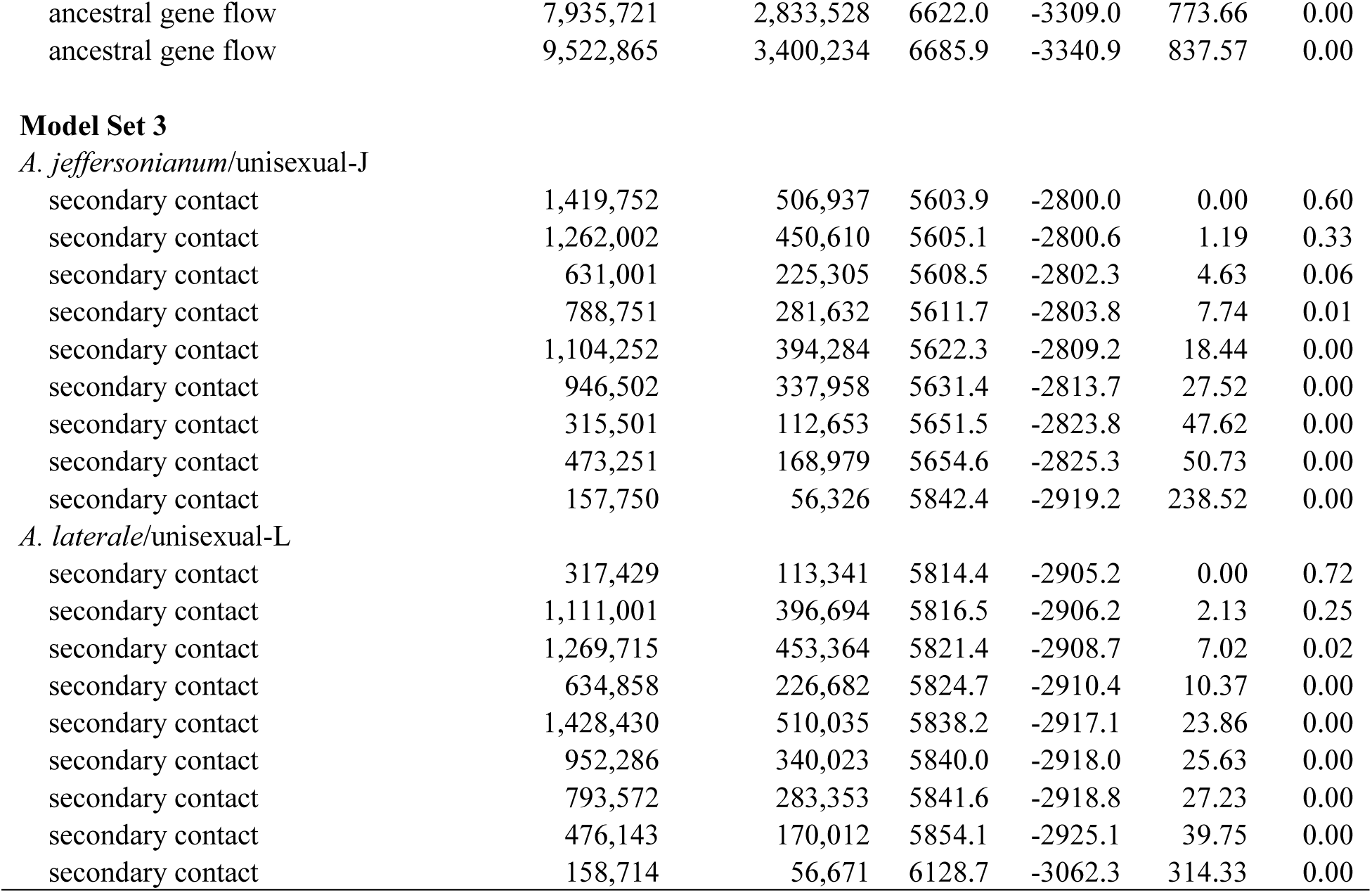
Results of PHRAPL model selection for both data sets that include *A. jeffersonianum* or *A. laterale* with their respective genomes recovered in unisexual individuals. The models sets are hierarchical: from five broad models (1), models including only secondary contact and divergence with gene flow (2), or models of only secondary contact (3). For 2 and 3, the time estimates for the end or beginning of gene flow are broken into equal increments from the present based on the best supported model from the previous set. Adjusted years are the estimates in years and scaled to the divergence date of the two sister species, *A. jeffersonianum* and *A. laterale* (12.4 mya). PHRAPL provides the Akaike information criterion (AIC), the log-likelihood (lnL), the change in AIC from the <u>top-ranked model (dAIC), and the weighted AIC value (wAIC).</u>

